# A common perceptual inference for cross-modally induced illusions of body schema

**DOI:** 10.1101/066159

**Authors:** Zane Z. Zheng, Kevin G. Munhall, Ingrid S. Johnsrude

## Abstract

Body-schema, or the multimodal representation of one’s own body attributes, has been demonstrated previously to be malleable. In the rubber-hand illusion (Botvinick & Cohen, 1998), synchronous visual and tactile stimulation cause a fake hand to be perceived as one’s own. Similarly, if a stranger’s voice is heard synchronously with one’s own vocal production, that voice comes to be attributed to oneself (Zheng et al., 2011). Multimodal illusions like these involve distorting body schema based on correlated input, yet the degree to which different instances of distortion are perceived within the same individuals has never been examined. Here we show that participants embraced the ownership of a fake hand and a stranger’s voice to a similar degree, controlling both for individual suggestibility and for general susceptibility to illusion of body schema. Our findings suggest that the perceptual inference that leads to the distortion of body schema is a stable trait.

## 1. Introduction

Our ability to make sense of incoming multisensory cues is crucial for the integrity of body schema. Mounting evidence has suggested, however, that body schema is plastic and subject to distortion as a result of perceptual inference about afferent multisensory signals (e.g., Botvinick & Cohen, 1998; Longo & Haggard, 2012; Maravita et al., 2003; Petkova & Ehrsson, 2008; Tsakiris, 2008; Zheng et al., 2011). Here we examine whether the malleability of body schema is correlated across two multimodal illusions that are both elicited through correlated input: the rubber-hand illusion (Botvinick & Cohen, 1998) which depends on correlated visual and tactile inputs, and the rubber-voice illusion (Zheng et al., 2011) which depends on correlated audio-somatosensory and audio-motor information.

In the rubber-hand illusion, *seeing* someone stroke a fake hand while *feeling* the stroking on one’s own hidden hand distorts hand ownership, producing the illusion that the fake hand is one’s own hand. The perceptual inference is that the visual input and the tactile input must be related because they are synchronous, and that, therefore, the fake hand must be one’s own. In the rubber-voice illusion, the vocal production of single words, when accompanied by temporally and phonetically matching auditory feedback in a stranger’s voice, elicits the illusion that the stranger’s voice is the talker’s own voice. This perceptual inference would seem consistent with everyday auditory experience where, when the act of speaking coincides with sensorimotor concomitants of speaking, the heard voice is normally attributed to one’s own.

It is clear that, despite the similar reliance on correlated sensory stimulation, the domains within which the rubber-hand and rubber-voice illusions are elicited are remarkably different (i.e., hand v.s. voice; visual-tactile v.s. audio-somatosensory and audio-motor). So does correlated input drive the perceptual inference of body schema similarly across domains? We believe that the two illusions provide a unique model where this question can be empirically studied, as the perceived strength of each illusion indexes the degree to which the product of the perceptual inference, i.e., distortion of body schema, is sensitive to the integration of coherent input. A correlation between the two illusions would suggest a perceptual commonality in the distortion of body schema, irrespective of domains.

To this end, we compare the two illusions within individuals, and conduct two procedures to exclude potential confounds. First, we test a uni-modal proprioceptive illusion in which stimulation of muscle spindles near the elbow causes the participant to perceive, inaccurately, that the forearm is moving (Lackner, 1988). This illusion is not contingent upon correlated multisensory input, and can therefore be used as a ‘baseline’ task to examine general susceptibility to illusion of body schema. We also administer the Multidimensional Iowa Suggestibility Scale (Kotov et al., 2007), a questionnaire that indexes an individual’s tendency to accept external suggestions. This would allow us to rule out a common factor related to the trait of suggestibility as a dominant explanation for any observed correlation between the multimodal illusions.

## 2. Results

The rubber-hand, rubber-voice, baseline illusions were assessed both subjectively and objectively. Subjective measures included questionnaire statements regarding the perceptual experience of distorted body schema. Objective measures were used to assess behavioral changes following the perceptual experience, in order to establish the validity of the illusion paradigms.

### 2.1. Objective and subjective measures of the three illusions

For the rubber-hand illusion, previous work indicated a post-induction displacement of the perceived finger position towards the rubber hand, as a result of induced rubber-hand ownership. Consistent with this finding, we also observed such a ‘proprioceptive drift’, where the perceived position of the participant’s left/hidden index finger shifted reliably towards that of the rubber hand after the synchronous stroking procedure (mean displacement ± SD: 2.95 ± 2.83cm; t(35) = 6.25, p < .001). The subjective ratings on the questionnaire statement assessing distortion of body schema (“It felt as if the rubber hand were my hand”) revealed a tendency to claim the ownership of the rubber hand (mean rating ± SD: 4.85 ± 1.57; t(35) = 3.25, p = .003, compared with a rating of ‘4’ on a 7-point Likert scale). There was no strong association between the magnitude of the drift and subjective ratings (see also Tsakiris & Haggard, 2005; Holle et al., 2011; Morgan et al., 2011).

For the rubber-voice illusion, we replicated the previous finding of ‘vocal following’ as the objective evidence for the development of the illusion (Zheng et al., 2011), such that participants were more likely to shift the fundamental frequency of their vocal production towards, than away from, that of the heard rubber-voice producing two stimulus words ‘day’ and ‘too’ (mean shift ± SD: for ‘day’, 15 ± 13Hz; for ‘too’, 15 ± 14Hz), χ^2^(1, N = 36) ≥ 5.444, p ≤ .02 (see Figure 1). Furthermore, participants’ perceptual experience of rubber-voice ownership (“It felt as if the voice I heard were my own voice”; mean rating ± SD: 3.85 ± 2.00 on a 7-point Likert scale) predicted the concomitant change in the acoustics (i.e., the shift of fundamental frequency) of their vocal production for ‘day’. (Spearman’s ρ = .398, p = .016; see Figure 2a). Note that this perceptually induced shift was not related to the acoustic difference between the fundamental frequency of the participant’s own voice and that of the rubber voice.

**Figure 1.**
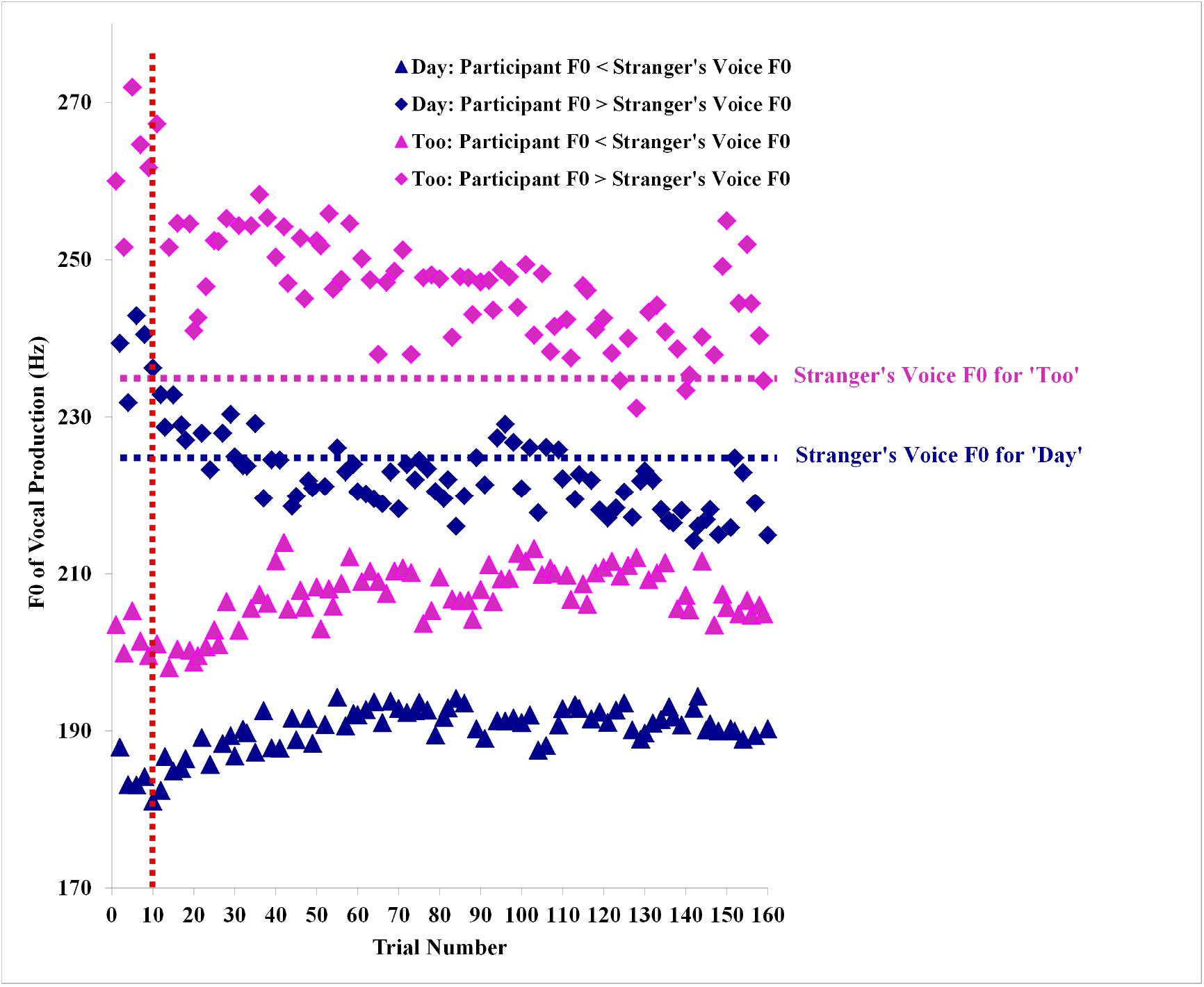
The converging patterns of participants’ fundamental frequency (F0s) that were higher and lower than the rubber-voice (or RV) F0 are separately shown for ‘day’ (in blue) and ‘too’ (in purple). The horizontal dash lines indicate the RV F0 for ‘day’ and ‘too’. The vertical dash line (in red) indicates the onset of RV auditory feedback.

**Figure 2.**
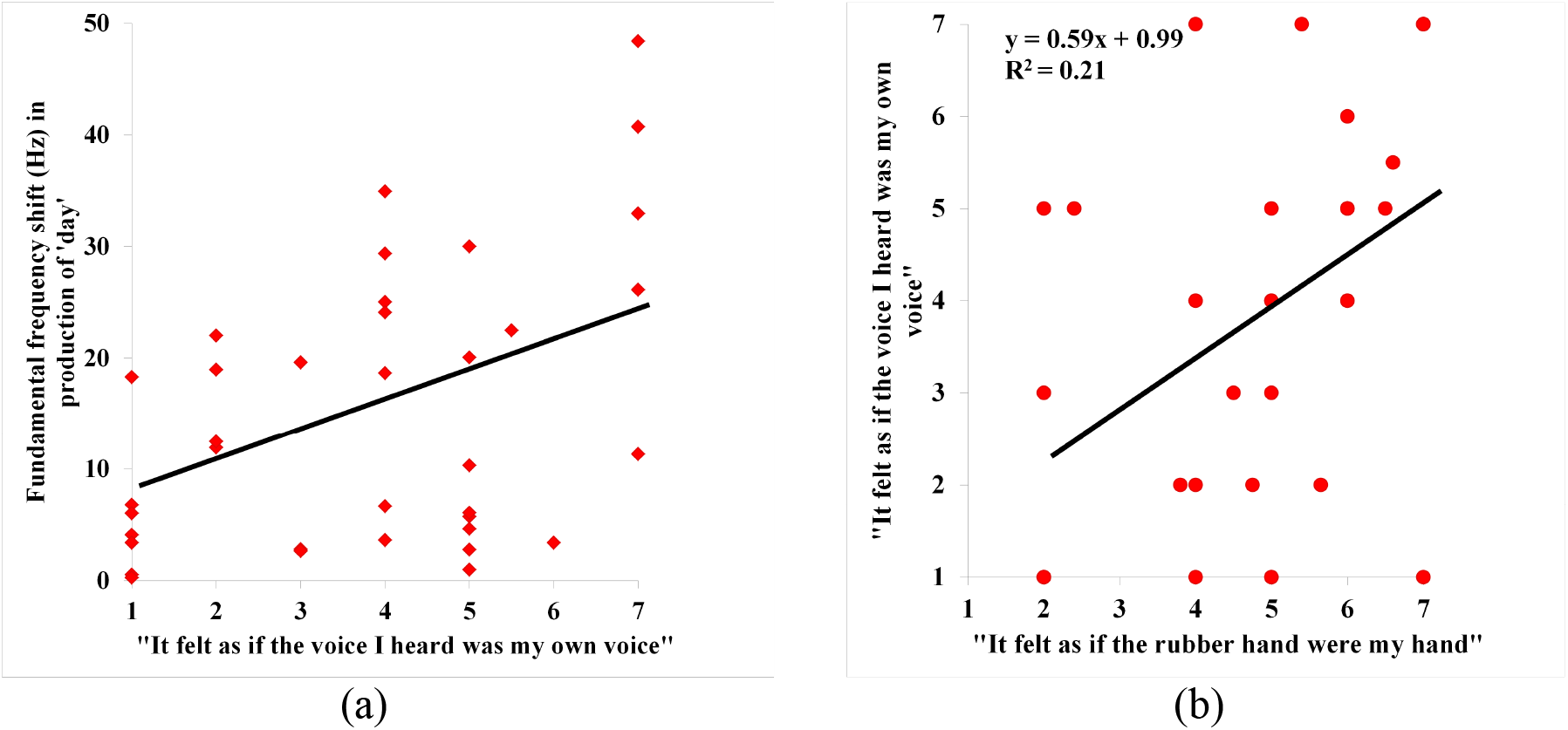
(a) The relationship between rubber-voice illusion ratings and shift in fundamental frequency (F0) is shown for ‘day’. The ratings were based on the question: ‘It felt as if the voice I heard were my own voice’ on a 7-point Likert scale. Participants who more strongly perceived the rubber voice as their own voice were also those who more strongly shifted their F0 during production. (b) Significant correlation (Spearman ρ = .510, p = .001) between the ratings of the rubber-hand and rubber-voice illusions is shown. Each illusion is represented by the key statement related to the distortion of body schema, i.e., “It felt as if the rubber hand were my hand” for the rubber-hand illusion and “It felt as if the voice I heard was my own voice” for the rubber-voice illusion.

For the baseline proprioceptive illusion, when asked to slowly move two forearms together with eyes closed, participants showed a strong left-right forearm positional asymmetry as a result of biceps stimulation (mean displacement ± SD: 2.66 ± 1.36cm; t(35) = 11.74, p < .001), with the stimulated forearm moving more slowly (as expected from a compensatory movement) than the unstimulated forearm. In general, they were negative about the distorted body schema statement (“It felt as if the forearm being vibrated were not my forearm”: mean rating ± SD: 3.08 ± 1.92; p = .007, compared with a rating of ‘4’ on a 7-point Likert scale).

### 2.2. Suggestibility Scale

To examine whether the perceptual effects of the three illusions might be predicted by individual suggestibility, we correlated participants’ scores on the Suggestibility Scale and their ratings on the body-schema statements of the three illusions. We observed a significant correlation between the scores on the Scale (mean score ± SD: 2.75 ± 0.49 on a 5-point Likert scale) and the ratings on the statement “I felt as if the forearm being vibrated were not my forearm” in the baseline proprioceptive illusion, Spearman ρ = .370, p = .026. This is not surprising, given the suggestive nature of the statement that is not directly linked to the perceptual effect (i.e., flexion) of physical stimulation and that is not driven by multimodal inference. In contrast, the Scale scores did not predict either the subjective or objective measures of the rubber-hand and rubber-voice illusions, suggesting that the induction of the two multimodal illusions was not simply due to individual suggestibility.

### 2.3. Perceived ownership of a rubber hand and a stranger’s voice predicts each other

We found that participant ratings on the body schema statements correlated significantly between the rubber-hand and rubber-voice illusions (Spearman ρ = .510, p = .001; see Figure 2b), but not between either of these and the baseline proprioceptive illusion, with the differences between the dependent correlations also being significant, p ≤ .006 (tested using the Meng et al., 1992 procedure). Thus, the degree to which participants perceived the rubber hand as their own hand predicts the degree to which they perceived the stranger’s voice as their own voice, but neither of these ratings predicted the strength of the baseline proprioceptive illusion. This lack of association with the baseline illusion ruled out the possibility that the strong correlation between the rubber-hand and rubber-voice illusions resulted from general susceptibility to body schema distortion.

## 3. Discussion

The rubber-hand and rubber-voice illusions emerge from integrating channels of multisensory information. Both illusions depend on the perceptual inference that synchronous, coherent activity across sensory channels must have a common origin (e.g., as predicted by Bayesian causal inference models; Kilteni et al., 2015), and both depend on distortion of body schema to make this inference possible. Our results demonstrate that the degree to which such synchronous, coherent activity across sensory channels drives perceptual changes in body schema is common across domains.

Previous work indicates that the ability to integrate sensory cues into a unified percept is highly variable (e.g., Ehrsson, 2012). Here we constrain this notion by showing that the ability to integrate multisensory cues for the perceptual representation of body schema can be stable within individuals, as indicated by the correlated perceptual ratings of the multimodal illusions. One might wonder why the objective measures were not correlated between the multimodal illusions. We note, however, that the objective evidence following the perceptual experience should be interpreted with caution. For the rubber-hand illusion, recent studies (e.g., Rohde et al., 2011; Abdulkarim & Ehrsson, 2015) have questioned a direct link between the proprioceptive shift and perceptual ownership of the hand, whereas for the rubber-voice illusion, the critical role of auditory/perceptual feedback in the control of vocal production behavior has long been established (e.g., see Brainard & Doupe, 2000). Given the distinct relationships between the perceptual and objective response, a lack of correlation between the objective measures of the two multimodal illusions does not necessarily weaken the strength of correlation observed for perceptual experience.

Our data does not support general susceptibility to body-schema distortion or individual suggestibility as the main explanation for the observed perceptual commonality. Here we offer two tentative interpretations that need to be elucidated in future studies. First, the perceptual commonality may reflect individual differences in the temporal binding of multisensory signals. Using a simultaneity judgment task, Stevenson et al. (2012) showed that individuals with more precise judgment in the synchrony of paired flash-beep stimuli were also more susceptible to the McGurk effect (McGurk & MacDonald, 1976). Given that the McGurk effect also critically depends on the integration of synchronous multisensory signals, it seems plausible to argue that what drives the shared hand and voice ownership here is associated with individual sensitivity to the temporal aspect of multisensory information. Alternatively, the operating mechanism underlying the perceptual commonality reflects a domain-general ability, modulating the degree to which multisensory integration drives perception of self-related attributes. If this is indeed the case, then partial ownership of body attributes might be more connected to the global/unified sense of selfhood than previously thought (e.g., see Blanke & Metzinger, 2009).

It is important to note that both rubber-hand and rubber-voice illusions are fundamentally ‘noisy’ (e.g., Shams, 2010): there are many parameters that can be variable. Despite the variability, we observed stability in distinct instances of body schema distortion. We believe that this main finding raises important questions that can be further explored. For example, are distinct body attributes uniformly represented at some level to facilitate the development of coherent self? And what is the role of multimodal integration in the inferential process that leads to the judgment of self when the decision space is less optimal?

## 4. Methods

### 4.1. Participants

Thirty-seven right-handed female students were tested. One participant was excluded for not following instructions in the rubber-voice procedure, leaving 36 (mean age ± SD: 21 ± 2). All participants were native English speakers and had no history of neurological/hearing impairment, and procedures were cleared by the Queen’s General Research Ethics Board.

### 4.2. Experimental Procedures

The three illusion-induction procedures, described below, were administered to each participant in random order, followed by the Suggestibility Scale.

#### 4.2.1. Rubber-Hand Illusion

The rubber-hand illusion was induced using a paradigm similar to that of Botvinick & Cohen (1998). Each participant was seated with her left arm resting under a mini-table (53.5cm × 30cm × 12cm), while a life-sized rubber model of a left hand was placed on the mini-table directly in front of her. During the experiment, a black blanket was used to cover the left forearm of the participant as well as a large portion of the mini-table, leaving the fingers of the rubber hand visible to the participant. The participant was instructed to fixate the rubber hand, while the experimenter used two identical paintbrushes to synchronously stroke the fingers of the rubber hand and the participant’s hidden hand, at a frequency of 1 Hz. The strokes were restricted to the middle and ring fingers to match the stimulus variety in the rubber-voice illusion procedure (see section 4.2.2.). What finger was stroked on each trial was pseudo-random, with the constraint that no more than three consecutive strokes were made on the same finger. There were 150 strokes in total.

Immediately before and after the stroking procedure, participants were required to indicate, with their right index finger, where they felt their left index finger (hidden under the mini-table) was located, as an objective measure of the illusion. With eyes closed, they drew their right index finger along the front edge of the mini-table until they judged it to be vertically aligned with the left index finger, which was resting directly beneath the mini-table during the stroking procedure. The horizontal displacement between the veridical position of the left index finger and its perceived position was measured using a standard ruler. At the end, participants were asked to complete a ratings questionnaire assessing the illusion on a 7-point Likert scale: ‘It felt as if the rubber hand were my hand’, ‘It seemed as if I were feeling the touch of the brush in the location where I saw the rubber hand touched’, ‘It seemed as if the touch I felt was caused by the brush touching the rubber hand’.

#### 4.2.2. Rubber-Voice Illusion

The rubber-voice procedure was documented previously (Zheng et al., 2011). Participants were seated in a single-walled sound-attenuating booth in front of a computer screen (Dell, Inc. U.S.A.). During the experiment, a word prompt appeared as text in the middle of the computer screen, once per second, and the participant immediately spoke this word into a microphone and heard concomitant auditory feedback through circumaural headphones (Sennheiser HD265 Linear, Sennheiser Electronic, Germany). The cue was one of two possible words (‘day’ or ‘too’), selected for being minimally variable in their acoustics across successive productions and across participants in a pilot sample.

The auditory feedback was processed and delivered by a real-time speech tracking system (iteration delay less than 10 msec; see Purcell & Munhall, 2006a,b) to ensure temporal and phonetic alignment with the participants’ own vocal production. Low-level white noise was present in the headphones to minimize bone-conducted speech feedback during vocalization (Barany, 1938). There were 10 trials of practice at the beginning during which participants alternated saying ‘day’ and ‘too’, and heard veridical auditory feedback. This was followed by a block of 150 trials, in which participants spoke the two words in a pseudorandom order (such that no more than three consecutive trials of the same word occurred) while hearing congruent, pre-recorded tokens of the two words in another female (i.e., stranger’s) voice. A previous pilot study demonstrated that each of 10 participants was able to distinguish her own voice from this stranger’s voice with 100% accuracy in a two-item forced-choice discrimination task.

The fundamental frequency of the participants’ vocal production was extracted across the 150 trials as a way of tracking the objective acoustic changes of produced vowels. At the end, participants were asked to complete a ratings questionnaire assessing the illusion on a 7-point Likert scale: ‘It felt as if the voice I heard was my own voice’, ‘It seemed as if the voice I heard was caused by my production of the word’.

#### 4.2.3. Proprioceptive Illusion

This baseline procedure was adapted from the proprioceptive illusion reported by Lackner (1988). Participants rested their forearm on the table and kept their eyes closed. During the experiment, they were instructed to slowly move their forearms up off from the table and toward their chest, while keeping the two forearms aligned with each other. The initial position and movement direction of the forearms were determined by considering previous work suggesting that participants perceive the illusion onset and respond faster when the forearms are held in a fully extended position at rest (e.g., Gooey et al., 2000). A hand-held electromagnetic physiotherapy vibrator (Touch ’n Tone, Conair Consumer Products Inc., Canada) was applied to the distal biceps tendon at the elbow on their left arm during the movement. The vibration stimulated the muscle spindles in the biceps that would otherwise cause muscle flexion, thereby creating a kinesthetic illusion that the forearm was moving. Consequently, participants compensated for the illusion by proactively moving the stimulated forearm in the opposite direction (in order to keep the two forearms aligned as instructed), leading to a displacement of the two forearms. When the right/unstimulated forearm reached a preset location (≈ 65^0^ angle between the forearm and the table), the participant was asked to end the movement, and the displacement of the two forearms was objectively measured as the horizontally projected distance between the knuckles of the middle fingers of the two hands. This procedure was repeated 3 times for each participant to obtain a stable estimate of the displacement. After the procedure, participants completed a questionnaire assessing the illusion on a 7-point Likert scale: ‘It felt as if the forearm being vibrated was not my forearm’, ‘It seemed as if I were experiencing forearm flexing at the time of the vibration’, ‘It seemed as if the flexing of my forearm was caused by the vibrator’.

In particular, the statement ‘It felt as if the forearm being vibrated was not my forearm’ was created (and phrased negatively) to match the perceptual consequences of the analogous body-schema statements in rubber-hand and rubber-voice illusions: e.g., just as stroking creates multisensory experience that may lead to illusory *de*-ownership of the real hand (i.e., ‘my real hand is *not* my hand’, see Longo et al., 2008) in the rubber-hand illusion, biceps stimulation in this proprioceptive illusion causes illusory movement of the forearm, which may lead to the experience of one’s own forearm not in control (i.e., my forearm *not* my forearm).

#### 4.2.4. Multidimensional Iowa Suggestibility Scale

The short Multidimensional Iowa Suggestibility Scale (short MISS; Kotov et al., 2007) was administered for each participant. The Scale used in our study contains 21 self-report items on a 5-point Likert scale. Ratings across the 21 items were averaged for each participant as an estimate of individual suggestibility (e.g., ‘When I see someone shiver, I often feel a chill myself’).

## Acknowledgements

This work was supported by a Lasell Faculty Research Grant (to Z.Z.), the Canadian Institutes of Health Research (to I.J.), and the Natural Sciences and Engineering Research Council of Canada (to KM and I.J.). I.J. was supported by the Canada Research Chairs program.

